# BehaveAI: a framework for rapidly detecting and classifying objects and behaviour from motion

**DOI:** 10.1101/2025.11.04.686536

**Authors:** Jolyon Troscianko, Thomas A. O’Shea-Wheller, James Galloway, Kevin J. Gaston

## Abstract

Here we introduce BehaveAI, a biologically inspired video analysis framework that integrates static and motion information through a novel colour-from-motion encoding strategy. This method translates object movement – direction, speed, and acceleration – into colour gradients, enabling both human annotators and pre-trained convolutional neural networks (CNNs) to infer motion patterns while retaining high-resolution spatial detail. Using a range of case studies, we demonstrate how the increased salience of motion information allows for the robust detection of objects that are challenging or impossible to identify reliably from static frames alone, particularly in complex natural scenes. We further demonstrate the reliable classification of different behaviours in animals and single-celled organisms. Additionally, the framework supports flexible hierarchical model structures that can separate the tasks of detection and classification for optimal efficiency, and provide individual tracking data that specifies *what* is present *where* and what it is *doing* in each frame. The framework makes use of the latest deep learning architecture (YOLO), combined with a semi-supervised annotation workflow. Together with salient motion information, these features can dramatically reduce the effort required for dataset annotation such that reliable models can often be made within an hour. Moreover, smaller annotation datasets mean that model training can be achieved on conventional computers without dedicated hardware, thereby improving accessibility. The motion encoding approach is also computationally lightweight, and can run in real-time on low-end edge devices such as a Raspberry Pi. We release the framework as a free, open source, and user-friendly package.

## Introduction

Videos convey rich information about *what* is present *where*, and crucially, what it is *doing*. This capacity to record spatiotemporal relationships and behaviour makes the use of video ubiquitous in society and indispensable for scientific research observing animal behaviour and ecological interactions. More widely, videos are used across sectors such as animal husbandry and farming, industrial quality control, security and surveillance, and medical diagnostics (Bouwmans, 2014). Nonetheless, despite decades of advancement in video processing, efficient and accurate quantification of complex motion information – critical for object detection and classification – remains a significant computational challenge, particularly in unconstrained visual scenes.

Convolutional neural networks (CNNs) have revolutionized the analysis of static visual information, with modern image classification, object detection, and segmentation tools demonstrating remarkable accuracy (Krizhevsky et al., 2012; Ren et al., 2016), and gaining widespread use in life sciences (Chan, Putra, et al., 2025; Thapa & Stachura, 2021), and wider society. Crucially, they can be trained effectively even with moderate-sized datasets due to techniques like transfer learning and data augmentation. These tools excel at recognizing objects based on their spatial features: shape, texture, colour, and pattern. Treating video analysis as a sequence of independent image processing tasks has yielded substantial progress (Chan, Putra, et al., 2025). However, this approach inherently neglects motion, which is critically important in many visual detection and classification tasks. For example, camouflaged animals can easily evade detection by blending in with their static surroundings, but they can become trivially easy to detect as soon as they move (Troscianko et al., 2009). Patterns of biological motion are also critical for recognising and classifying behaviour, as illustrated by the relative movements of a few dots that are instantly recognised as a human walking (Johansson, 1973).

Tools that integrate motion information for detection and classification remain less widespread than static variants. Examples include DeepEthogram, which can effectively determine patterns of animal behaviour through static and motion information (Bohnslav et al., 2021), although the tool is not suited to the detection, classification and tracking of animals in typical natural scenes. LabGym detects temporal changes in animal outline shape to predict its behaviour, and supports tracking the behaviour of multiple individuals (Hu et al., 2023), however the model training process is complex, particularly when detecting objects against natural backgrounds. Pose estimation tools such as DeepLabCut and SLEAP are effective for determining the spatio-temporal movements of whole animals and their limbs (Mathis et al., 2018; Pereira et al., 2022), but these data don’t immediately tell us what the animal is doing without considerable further interpretation (Hardin & Schlupp, 2022). Tools such as Keypoint-MoSeq can use pose for behavioural classification (Weinreb et al., 2024), however, training workflows for pose estimation are typically time consuming, limiting their accessibility. Critically, tools that rely on outline shape or pose fail when bodies and limbs cannot easily be resolved, e.g. due to small limbs and video resolution limits, motion blur, fast movement, complex backgrounds, occlusion, or limb transposition (the limbs of nearby individuals causing confusion). These are common circumstances in real-world videos, particularly considering that most animals have evolved colouration, movement, and background selection in order to evade detection. 3D CNNs such as ConvNet can combine static and optic flow streams for effective video classification (Carreira & Zisserman, 2017; Simonyan & Zisserman, 2014), although this system is not readily suited to object classification and tracking tasks. Notably, these video analysis tools have not received widespread use in behavioural and ecological sciences when compared to static CNN frameworks despite the fields’ heavy reliance on video data (Hardin & Schlupp, 2022) and common need to quantify behaviour.

The mammalian visual system uses segregated neural pathways for different visual tasks. The ventral stream primarily processes form, colour, and object identity – the ‘what’ of the scene. The dorsal stream deals with motion, spatial relationships, and guiding action – the ‘where’ and ‘how’ (Milner & Goodale, 2006, Chapter 1.3.3). These functionally distinct processing streams are then integrated to create a complete percept. Inspired by this segregated but integrated processing, we present a novel framework specifically designed for robust object detection, behavioural classification, and individual tracking from video data. Our framework converts motion information into false colours that allow both the human annotator and FCNN easily to identify patterns of movement (fig. 1). Distinct from motion history images (Komori et al., 2023), outline shape changes (Hu et al., 2023) or optical flow based kinematics (Carreira & Zisserman, 2017; Komori et al., 2023), our colour-from-motion strategy provides information on the direction, speed, and acceleration of movement through colour gradients, with different colours reaching further back in time. This motion encoding method uses a trivial amount of computational power and shifts the motion interpretation to the pre-trained FCNN while also retaining high-resolution spatial information, meaning that the framework can achieve real-time video processing on low-end ‘edge’ devices. The system can also integrate conventional static (colour, pattern, form) information together with hierarchical processing, meaning that the tasks of detection and classification can use motion and/or static visual information, much like the mammalian visual system. We further introduce a semi-supervised annotation workflow for rapid and efficient ‘auto-annotation’.

**Figure 1.**
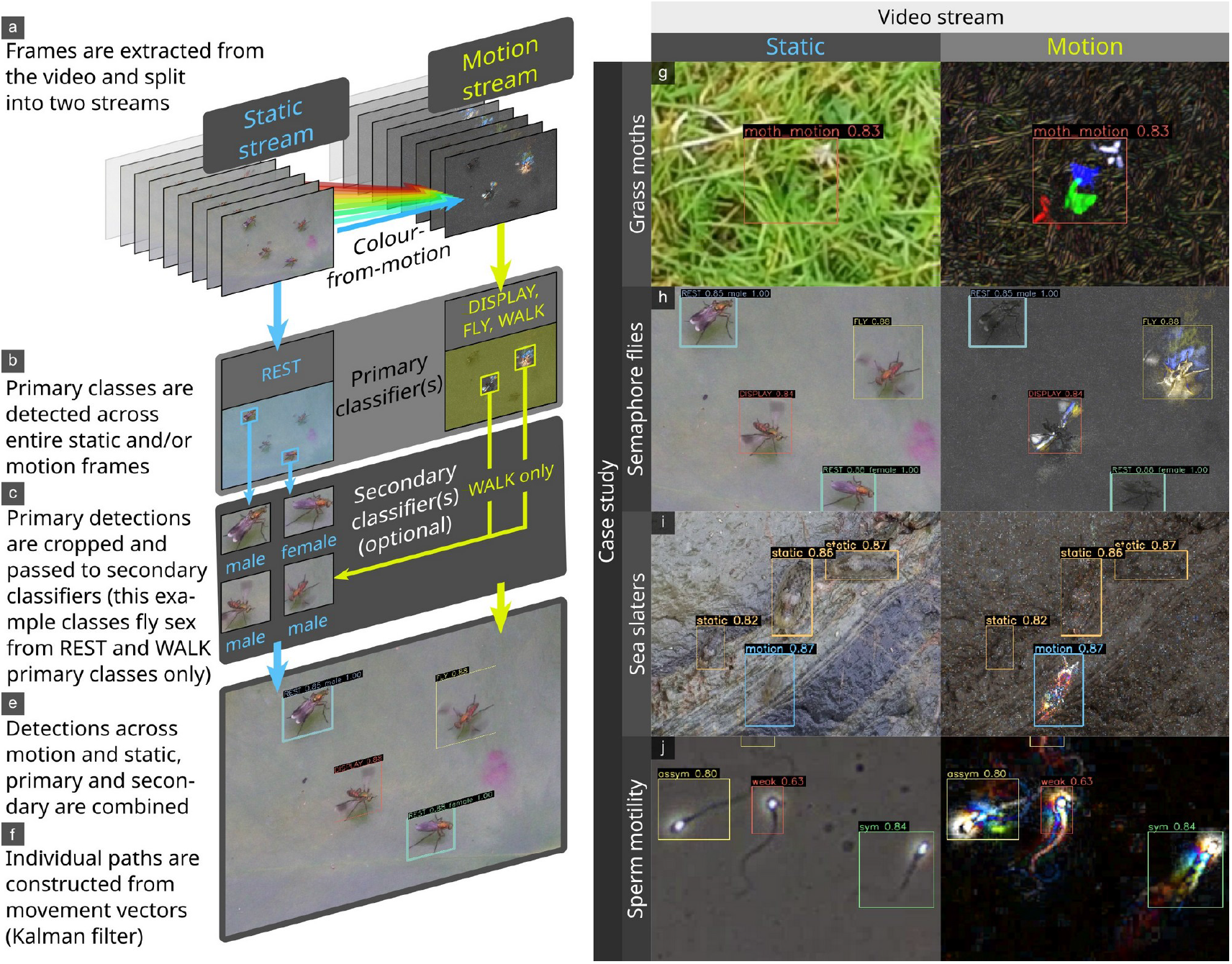
Illustration of the framework’s structure (a-f). Users can specify primary classes and secondary classes across both static and motion video streams. The right panel shows examples of static and colour-from-motion frames from case studies. The colour-from-motion trails create characteristic patterns that reveal movement and behaviour. The grass moth (Crambidae) is nearly undetectable from still frames due to its colour, and motion blur, but it is highly salient in the motion stream (g). The semaphore fly (*Poecilobothrus nobilitatus*) example shows how motion information can easily disentangle behaviours that are often identical from static frames (e.g. ‘fly’ and ‘display’ appear identical in the static frame, but different in motion). This example also showcases hierarchical classification, with secondary classifiers determining the sex of the flies (h). Sea slaters (*Ligia oceanica*) are highly camouflaged when static, and salient when moving, resulting in motion models that make far fewer errors (but cannot detect stationary individuals) (i). Human sperm have been classified based on their swimming movement with either symmetric (typically resulting in fast, straight movement), asymmetric (typically resulting in slow, circling, exploratory movement), or weak (twitching, vibrating etc…) strategies. These swimming strategies can be determined without tracking individuals, which is difficult in complex, debris-filled videos (j).

Here we describe the framework and demonstrate its ease-of-use and efficacy in a wide range of challenging object detection and classification tasks. Although we provide some comparisons, we have not focused on directly equivalent benchmarks between our methods and others due to the dangers of attempting to over-generalise AI model utility (Raji et al., 2021). Indeed the FCNN architecture that our framework relies on has been demonstrated elsewhere (Jocher et al., 2023), thus we instead outline BehaveAI’s potential to transform its capabilities. Direct benchmarking presents further practical limitations: i) existing annotation sets would all require re-annotation for the motion stream, but ii) we aim to show that comparable performance can be achieved with much lower user annotation effort; and iii) we note that many existing datasets do not provide videos that match typical natural scenarios (complex, moving backgrounds, variable lighting), and often select behaviours based on convenience and visibility rather than end-user requirements.

## Results

Our case studies test the performance of the BehaveAI framework in a range of challenging tasks, and we use these to outline key advances. First, movement is typically far more salient than static visual cues, increasing the signal-to-noise ratio of objects against their backgrounds, and making the models more independent of background appearance or lighting conditions (figs. 1-3). The grass moth case study shows how a motion-based model can significantly outperform static-based models for equal annotation effort, with precision and recall of 0.966 and 0.969 (motion, 98.3% correct validation inferences, i.e. 115 correct detections for 2 false negatives) versus 0.792 and 0.544 (static, 67.5% correct validation inferences). We give the precision and recall metrics as provided in the final epoch of model training by YOLO. Precision indicates how many detections were correct, and high numbers imply few false-positives. Recall reflects the model’s ability to identify all instances of objects, and high numbers imply that few objects are missed. The pigeon case study similarly achieves comparable performance to Chan *et al*. (2025) who used 1,587 training images, while we used just 295, despite us adding an additional behaviour class (see fig. 4). The sperm motility case study can classify subtle differences in sperm movement with precision 0.793, and recall 0.801 based on 333 annotated motion frames, while using static frames alone reduced model performance to 0.610 and 0.594 respectively. Other studies show that classifying sperm defects from static frames is considerably more difficult, for example, Thambawita *et al*. (2023) use annotations from over 29,000 frames to achieve precision 0.571 and recall 0.228 with the same dataset.

A second major benefit relates to the fact that most animal (and single-cell/subcellular) behaviour is characterised by biological motion, and by training the model directly on the motion cues – not just speed and direction, but also spatial patterns of acceleration and deceleration – we can bypass the complex and computationally demanding steps required to predict behaviour from pose estimation or whole-individual-tracking. This is highlighted by the sperm motility example (above), where tracking the progress of individual sperm (which is difficult in when multiple individuals crowd together) is not required in order to assess the frequencies of different sperm motility types in a given sample (figs.1, 4 & 5).

Third, behaviour can be classified from statically-identical frames, as highlighted by the pigeon example, where a motionless pigeon often has an identical pose to a preening or walking individual (figs. 1 & 4). Similarly, in the semaphore fly example, a flying insect can appear identical to a courtship displaying insect, and a walking insect is visually identical to a resting insect in a static frame (fig. 1h). In this latter case, incorporation of information allows behaviours to be classified with precision 0.895 and recall 0.894 (correct inferences for display: 97%, fly: 94% and walk: 94%).

Fourth, movement can often be detected and classified from a far lower spatial resolution than static classifiers. The grass moth scaling test (fig. 3) shows how the motion-based model can perform very well: precision and recall of 0.945 and 0.949 respectively, (>96% correct validation inferences) when the static targets’ bounding boxes are just 4.3 pixels across on average, and this drops to just 0.813 and 0.795 (82% correct validation inferences) when targets are just 2.0 pixels across on average. Static model performance at these scales is 0.633 and 0.538 (36.4% correct) and 0.225 and 0.3117 (27.3% correct) respectively.

Finally, the framework’s colour-from-motion strategy is extremely computationally efficient, enabling real-time detection and classification on low-end devices. We demonstrate the framework running in real-time on a basic Raspberry Pi 5 at 8-9 fps with inference being performed on input video resolution of 640×480 pixels.

## Discussion

The BehaveAI framework combines a novel colour-from-motion strategy with state-of-the-art deep learning architecture in an integrated pipeline. Our results showcase the accuracy, flexibility, ease-of-use, and computational efficiency of the framework for detecting and classifying objects from their movement. Specifically, we show how the framework can achieve comparable or greater inference performance from far smaller annotation training sets than alternative methods, and is able to track objects that might not be possible with other methods. This in turn brings further benefits: reducing training time, allowing models to be trained and deployed on low-end computers, simplifying analysis, and reducing energy consumption. This also eliminates the need to purchase cloud-based compute services, reducing costs and barriers to entry. Notably, the framework also excels at tracking small, camouflaged moving objects – just two pixels across – that cannot be reliably detected from still frames. This means that a motion-based strategy could effectively cover a much larger spatial area and still detect targets with far greater accuracy than a static model. Alternatively, it shows how video streams can be reduced in size substantially without affecting performance, thus increasing computational efficiency.

Determining *who* is doing *what, where*, and *when* will often require a more complex set of classification tasks than can easily be achieved in a single classifier. The BehaveAI framework’s biologically inspired separation of motion and static video streams, together with hierarchical processing allows for considerable flexibility in structuring the pipeline to meet individual needs. This is highlighted in the semaphore fly example, where flies are first detected by primary models that search across whole frames for specific motion (walking, flying, displaying), or static appearances (rest). Once detected by a primary model, the flies are passed to a secondary model that determines the sex of the fly from the cropped region, increasing computational efficiency and reducing redundancy. This maximises the efficacy of the initial detection task – whether flying quickly or completely still – and only then determines the sex based on subtle visual cues.

Manual annotation of objects in images/videos is a crucial bottleneck in the deployment of deep learning models, typically requiring considerable time investment before performance can be assessed. The sheer diversity of systems, organisms, and behaviours observed within the biological sciences means that generalised ‘foundation’ models often fail to transfer between systems, requiring researchers to develop bespoke models for their specific use cases (Chan, Brookes, et al., 2025). Efforts have therefore focused on optimising this process, rather than attempting to build generalised datasets (Beery et al., 2018; Raji et al., 2021). The BehaveAI framework uses a semi-automated annotation workflow, which – combined with the above increase in salience from motion cues – results in a far more efficient annotation process. By utilising model predictions to augment annotation, the number of manually drawn annotations required is considerably reduced, while cases where the model performs poorly are highlighted, enabling the user to target additional data for inclusion only as needed. As above, this reduces the volume of data needed to develop a robust model and limits training and deployment costs downstream. The grass moth example (figs. 1-2) details how – within one hour – roughly 550 frames can be annotated and a model trained that achieves remarkable tracking performance. This is despite the videos having substantial background motion due to grass swaying in the wind and the camera moving. We also include tools for inspecting the annotation dataset (in order to fix errors or re-classify annotations), and for rebuilding the motion annotations in order to compare the efficacy of different colour-from-motion strategies. Together this makes for a user-friendly workflow that maximises efficiency and accessibility.

Implementing edge BehaveAI models on low end hardware opens up a range of opportunities for real-time monitoring and interaction with behaviour in the field or lab. We demonstrate that the framework’s inference processes can run on Raspberry Pi 5 units, with YOLO11 models converted to NCNN format for computational efficiency. The increased salience of motion information means that video frames can be reduced in size substantially, in order to increase speed, without impacting performance. This demonstration highlights the versatility of the BehaveAI framework for deployment in embedded and remote scenarios.

The current BehaveAI framework provides unique opportunities for efficient and robust behavioural quantification, yet there are a number of avenues to explore for future enhancement. Integrating existing annotated datasets is not currently straightforward because the boundaries for motion annotations will differ from the static; ongoing development will aim retroactively to extract motion information from such data, although this would only be possible for datasets where full videos with annotations are available. Beyond this, we aim to incorporate support for segmentation, pose estimation, and orientated bounding-box annotations. Together with these enhancements, the open source code and documentation provided here will enable the incorporation of our motion strategy into a wide range of aligned deep learning frameworks. The utility of the framework in fields beyond the biosciences should also be explored, for example, its ability to quantify acceleration and deceleration of objects within a single frame could be valuable in machine vision tasks such as autonomous vehicle control.

We aim to make the BehaveAI framework as accessible as possible, and release the tools as a free and open source software package. We also include a range of tutorial videos and sample data. The framework’s flexible structure means that switching the pipeline to use alternative deep learning architectures is straightforward, allowing the system to upgrade easily to keep pace with the latest advances in deep learning.

## Data availability & Online resources

Software download: https://github.com/troscianko/BehaveAI

Introduction video: https://www.youtube.com/watch?v=YQG4497kzPY

User guide video: https://www.youtube.com/watch?v=PiX7Fp2F-Xk

Case study data: DOI:10.6084/m9.figshare.30531116

## Case Studies

### Grass moths

Our grass moth dataset presents a challenging but common detection task in ecology: tracking small animals that are motion-blurred and often occupy just a few pixels, travelling fast against complex, moving backgrounds with an abundance of wind-blown vegetation distractors of a similar size, colour and shape. In any given static frame the moths are barely visible, but their motion is typically highly salient (figs. 1-2). The dataset is made up of 32 short video clips of individual grass moths (Crambidae, unknown species) taking off as they are approached on foot in a grazed pasture field in the UK with varied lighting conditions (sunny to cloud-covered). Filmed using a hand-held smartphone (Huawei p30, 1920×1080, 60 fps [frames-per-second]). We used the ‘sequential’ motion strategy with a YOLO11n model, 50 epochs per training round (full settings and data are available in the repository [LINK]).

First, we use the dataset to demonstrate the speed with which an effective moth tracking model can be built. Training initially used 77 annotated frames from 6 videos (38 additional annotation frames were automatically set aside for validation), after which the first model was trained. Following this, semi-supervised annotation (‘auto-annotation’) was used to add another 168 training annotation frames from 7 videos (plus 52 set aside for validation), and a second model was trained. At this point auto-annotation was making very few errors, and the final round of annotation focused on correcting and adding blank frames that had been auto-annotated with false positives, amending false negatives, and adding low-confidence auto-annotated hits, yielding another 194 training annotation frames (plus 43 set aside for validation) from 6 new videos. Only a small fraction of these annotations were manually drawn. This entire annotation process (548 annotated frames, of which 57 were blank frames, including two rounds of model training), took 61 minutes to complete. The confusion matrix of the final model shows that 114 moths were correctly detected in the validation set (98.3%), with 2 (1.7%) false negatives, and 7 false positives; precision: 0.966; recall: 0.977. Note that these false positives would be easy to exclude post-processing by filtering out any short tracking paths based on tracking ID.

In order directly to compare the performance of motion-based classification to a conventional static classifier we annotated the same grass moth videos from the static stream. Given the moths’ speed, the motion annotations must be drawn over a considerably larger area than the static ones, so separate annotations were made. The moths are often not distinguishable from their background in static frames (e.g. wings folded up or down, with motion blur and similar colour and shape to background features, see fig. 2), and were only included if they were visible (meaning the static model could never match the motion model’s ability to detect the moths in all frames). The motion dataset used 415 frames for training and 133 for validation, from 19 videos; the static dataset used 460 frames for training and 103 for validation. YOLO11s models with 150 epochs were used for training. The motion model achieved precision and recall of 0.966 and 0.969 respectively (fig S1; of 115 correct classifications, there were 2 false negatives and 14 false positives). The static model achieved precision and recall of 0.792 and 0.544 respectively (fig S2; of 52 correct classifications there were 25 false negatives and 32 false positives). But as noted above, this over-estimates practical performance of the static model as the moth was not distinguishable from its surrounds in a large proportion of frames, making continuous tracking from static frames far more problematic than from motion.

**Figure 2.**
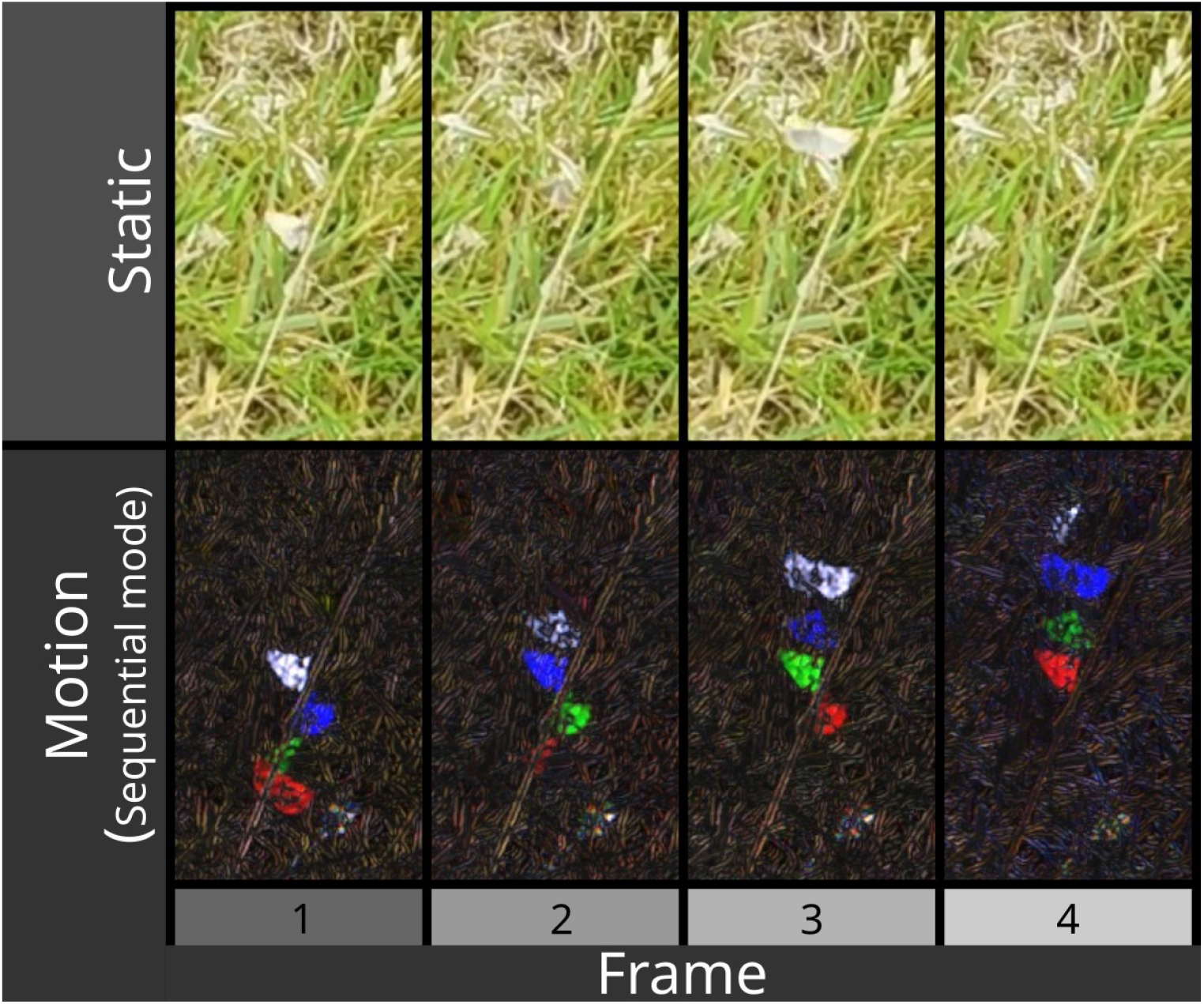
showing a grass moth (Crambinae) in flight, cropped over consecutive frames. The moth is barely visible in static frames (top), but becomes highly visible in motion (using sequential mode in this case).

Next, we tested how target size affects model performance. Images in the above training sets were reduced in resolution to 1/2, 1/4, 1/8, and 1/16 (0.5, 0.25, 0.12 and 0.06) of their original scale, and models were re-trained using the same settings. Rescaling was performed using the batch conversion tool in ImageJ v1.52, with bicubic interpolation. The results are shown in fig. 3. The motion model’s precision and recall were greater than 0.945 and 0.949 respectively down to 1/8th scale (image dimensions 240×135 px, target dimensions 4.27 px), and reduced to 0.813 and 0.795 at 1/16th scale (image 115×64 px, target 2.04 px). The static classifier performed worse, at all scales, with precision and recall greater than 0.633 and 0.538 down to 1/8th scale, and 0.225 and 0.312 at 1/16th scale

**Figure 3.**
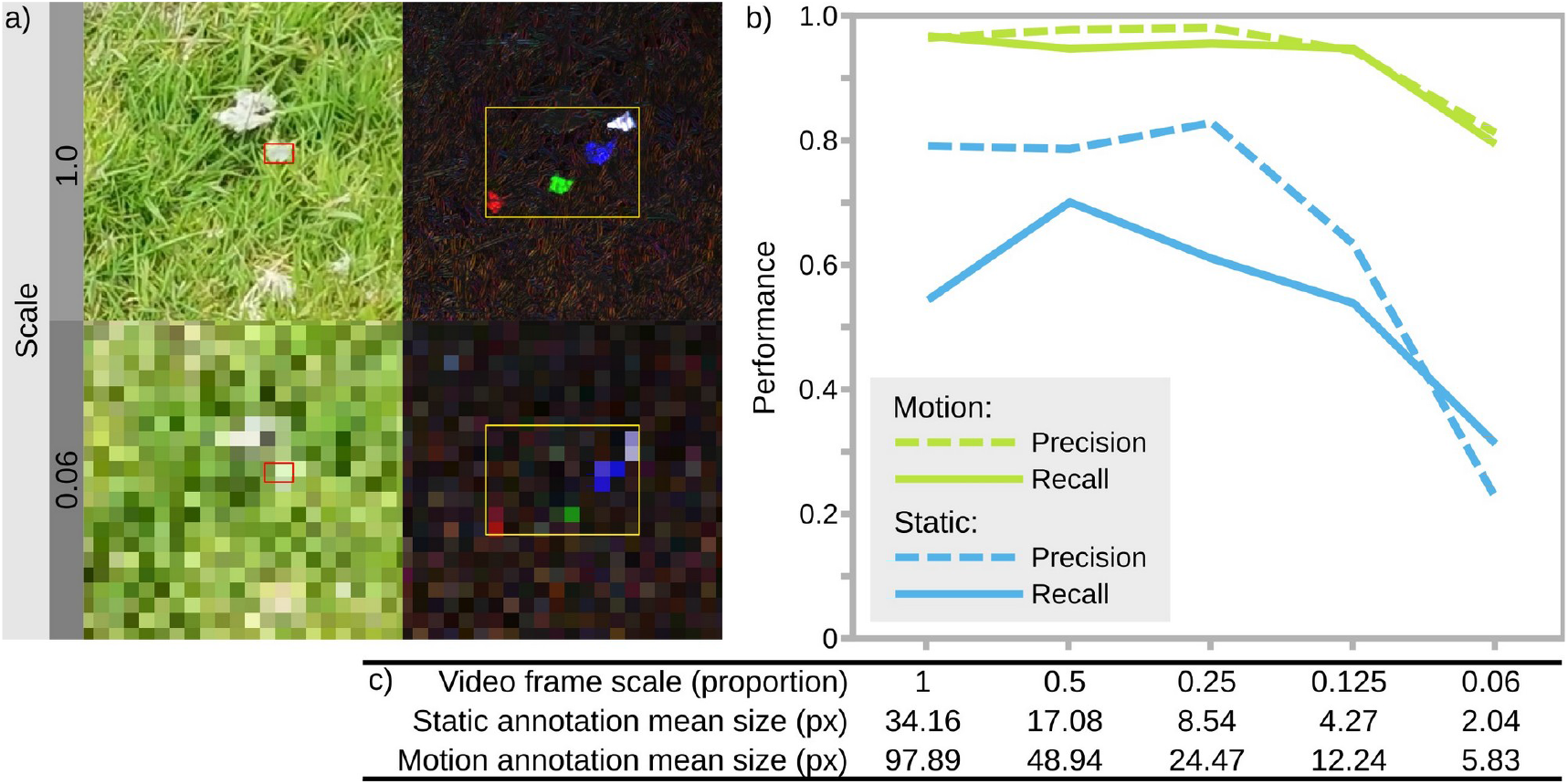
Motion and static classifier performance across different target sizes. Grass moth annotation images were progressively scaled down, and models re-trained on each resolution. a) shows cropped regions with unscaled (1.0) and 1/16th scaled examples, with boxes showing the bounds of the static (red) and motion (yellow) grass moths. Movement creates a substantially larger and more salient target for detection. b) shows model metrics at different scales; the motion model performs consistently high at all scales, but the static model performance decreases markedly with scale. The performance of the static model is over-estimated here because moths are simply not visible in a substantial number of frames. c) lists the scales and respective mean annotation box sizes (in pixels). Videos are 1920×1080 pixels at scale = 1.0.

### Pigeon behaviour – 3D-POP dataset

We used a subset of the publicly available pigeon behaviour dataset [3D-POP, (Naik et al., 2023)] for comparing performance with and without motion strategies. We annotated 295 training images, and 70 validation images, taken from 4 training videos (considerably fewer than Chan et al. 2025, who used 1,587 training images). We specified behaviour classes similar to those of the original publication and those used for benchmarking by Chan et al. (Chan, Putra, et al., 2025). However, we used the broader definition of ‘courtship’ to refer to all male courtship behaviour, rather than the ‘bow’ alone now that we are not constrained to cross-verify with 3D body-tracking data. We also added a ‘still’ class for motionless birds. Given that the birds were slow-moving against a uniform background, annotation boxes drawn for motion were also suitable for static frames, allowing for direct comparisons without redrawing annotations. Three classifiers were trained from the same annotation data by altering the image processing strategy; ‘static’ (conventional motionless frames, fig. 4a), ‘motion (default)’ (exponential motion strategy, fig. 4b), and ‘motion (chromatic tails only)’ (exponential strategy with chromatic tails only set to true, fig. 4c). Each was trained with 100 epochs using YOLO11s. Confusion matrices for each strategy are shown in fig. 4. The static model achieved precision and recall of 0.651 and 0.677, the default motion model achieved 0.788 and 0.780, and the motion-tails-only model achieved 0.746 and 0.787 respectively.

**Figure 4.**
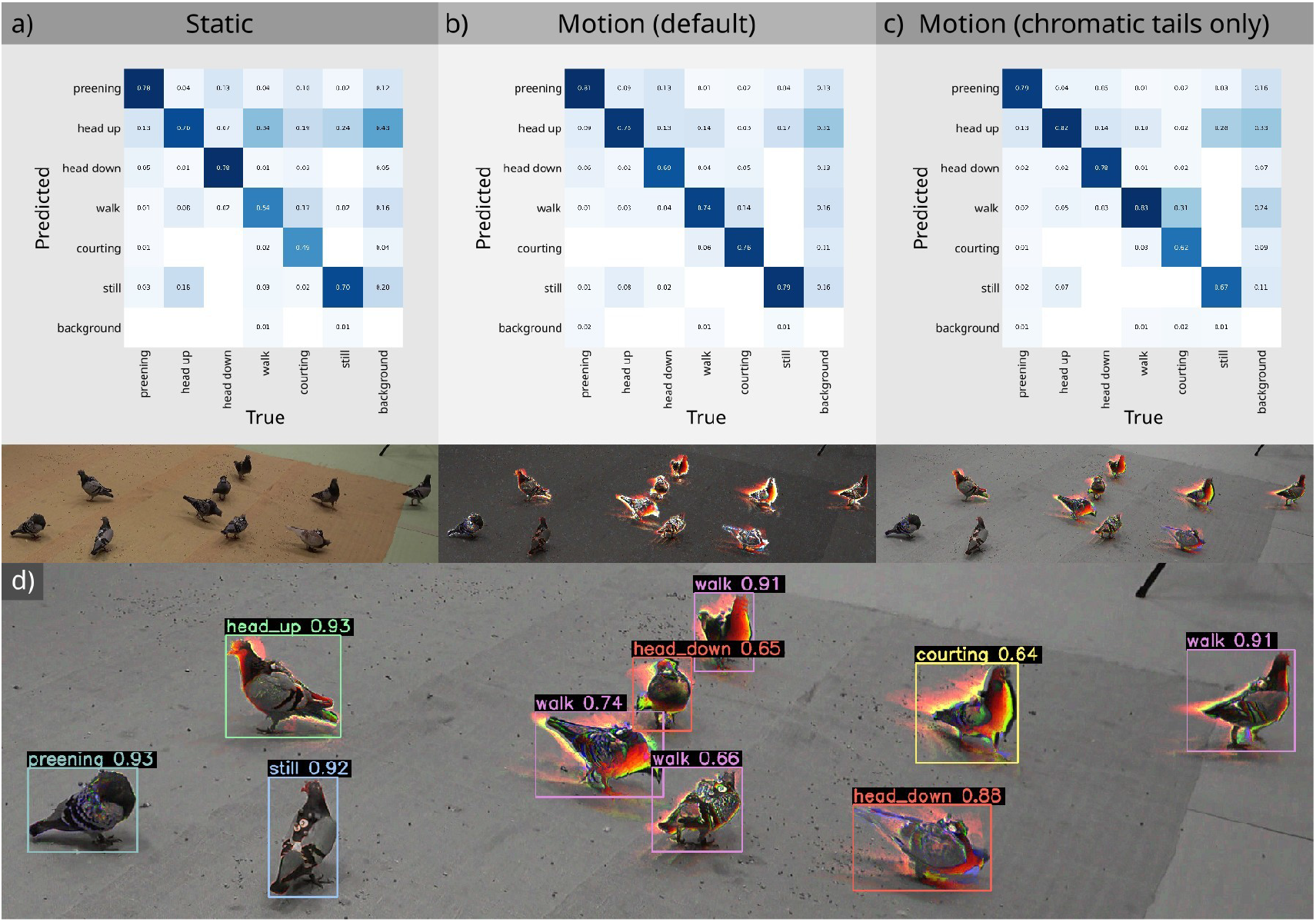
Direct comparison of pigeon behavioural classification using static frames (a) and two different motion strategies (b-c). (d) shows an auto-annotated frame using an exponential strategy with chromatic tails only, correctly classifying all of the pigeons and their behaviours. Numbers show the auto-annotation confidence.

### Sperm motility classification

Quantifying sperm motility is important for assessing human fertility, and a range of tools have been developed to automate the process. Nevertheless, the task remains difficult, particularly given the wide range of imaging systems, tail movements that are often difficult to track, and samples with a considerable degree of debris and visual noise. We used the VISEM video dataset of human sperm (Thambawita et al., 2023) for training a motion classifier to distinguish between three characteristic sperm motion strategies; symmetric, asymmetric, and weak motion (see fig. 5 for examples). The training set consisted of annotations from 333 frames using 16 videos, plus 74 validation frames. A YOLO11m model was trained with 200 epochs, resulting in a precision 0.793, and recall of 0.801 (fig. S3). A key advance here is that the motion behaviour of a sample can be reliably ascertained from a single frame without needing to track the movement of specific individuals (which is problematic in complex scenes).

**Figure 5.**
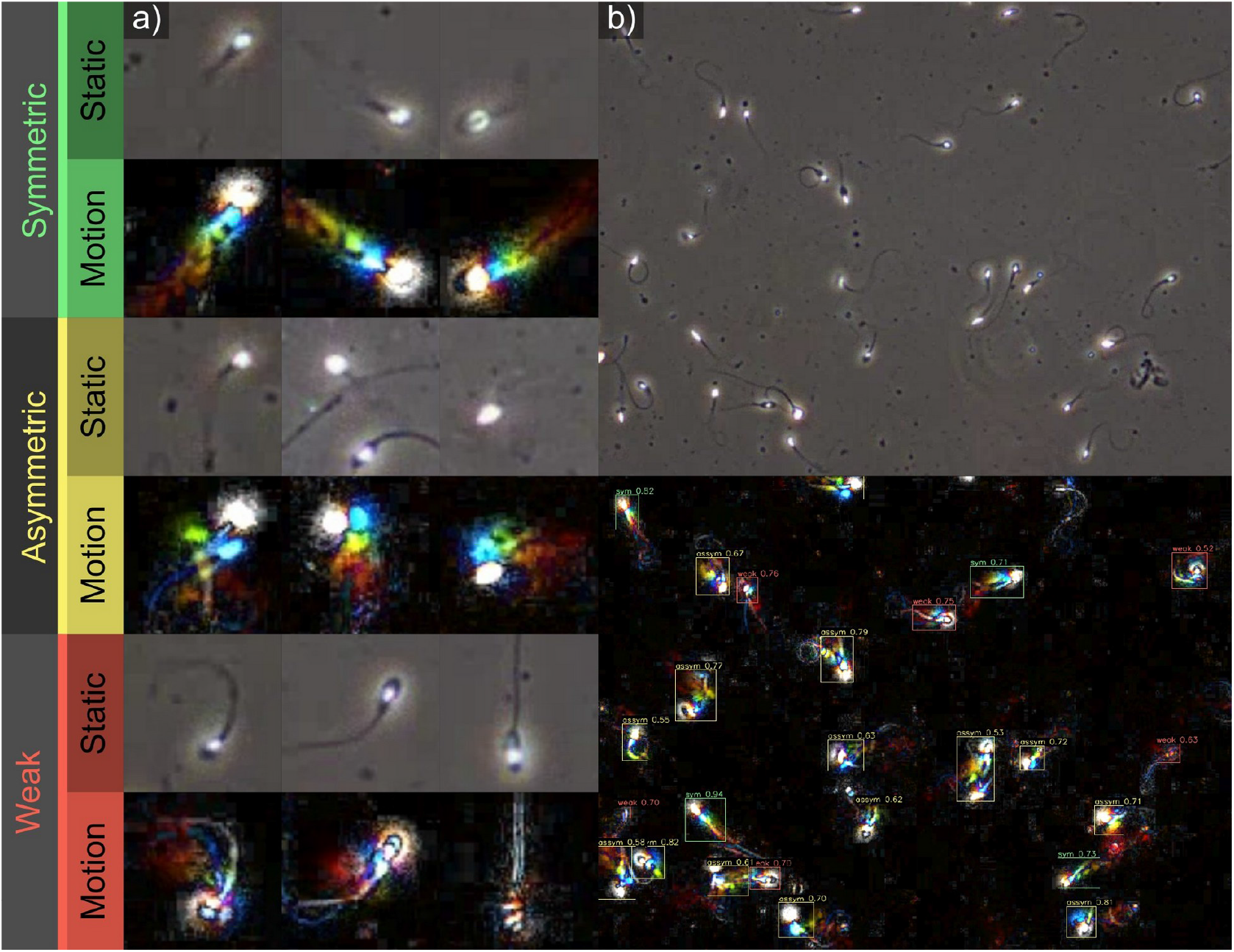
a) Shows examples of static and motion (exponential mode) streams for the three types of motion being classified. Symmetric movement results in a linear, consistent motion trail behind the head. Asymmetric motion results in a zig-zag, irregular motion trail, while weak motion (vibrating/twitching sperm) creates motion energy without a clear trail. b) shows a full-frame example from the dataset with classifications and confidence shown in the bottom panel.

**Figure 6.**
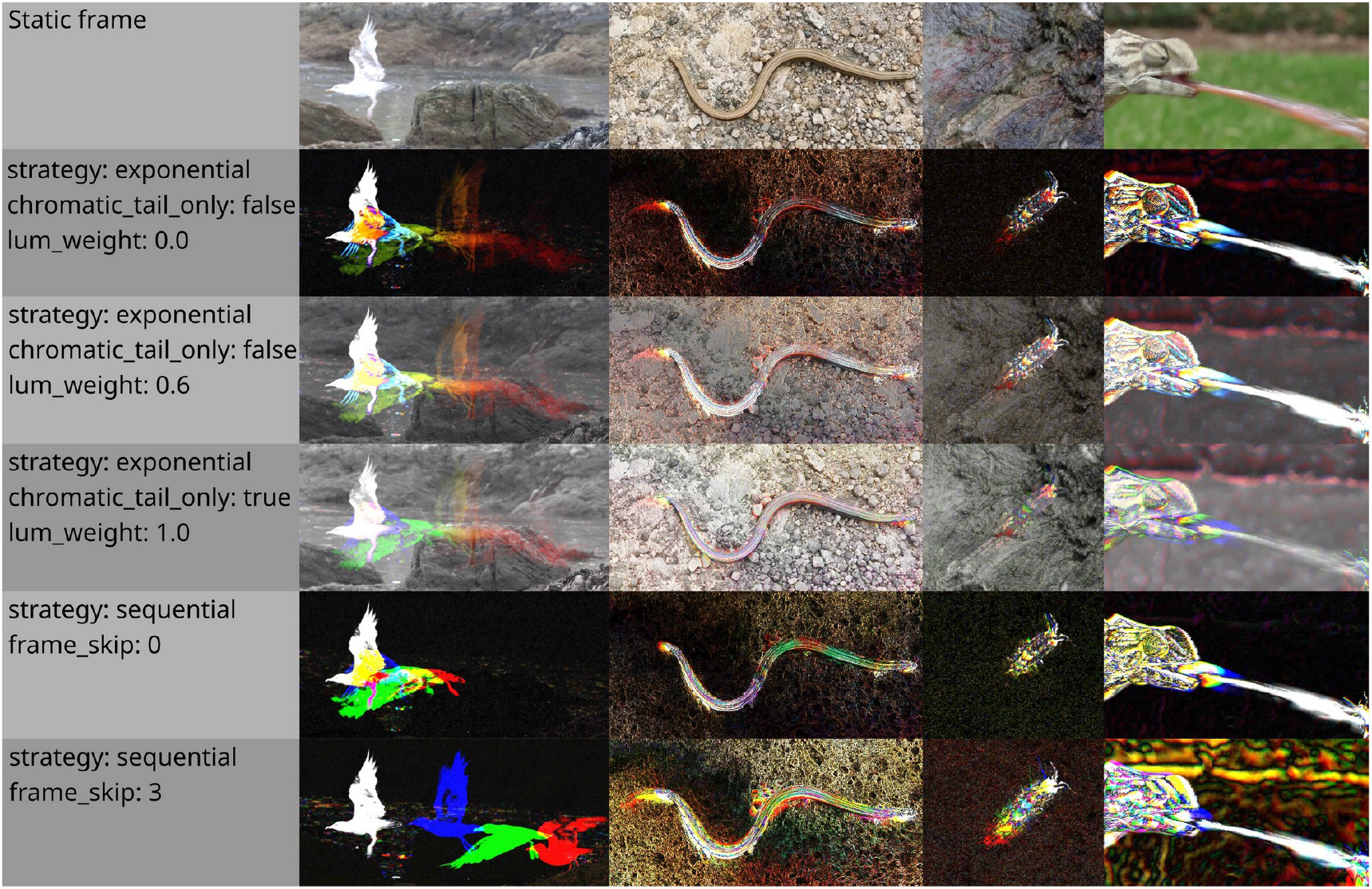
showing a selection of colour-from-motion strategies and parameters available in the BehaveAI framework. Examples from left to right are a herring gull (*Larus argentatus*) taking flight, a common slow-worm (*Anguis fragilis*), a camouflaged sea slater (*Ligia oceanica*) moving, and a flat necked chameleon (*Chamaeleo dilepis*) hunting an insect. The motion-coding parameters can be altered after building the annotation training set, and new models trained, allowing users to test which parameters best suit their system without redoing annotations.

Training the model using static frames alone (and all other parameters kept the same) yielded precision and recall of 0.610 and 0.594 respectively (fig. S4).

### Sea slater detection

Sea slaters (*Ligia oceanica*) are littoral-zone isopods that exhibit polyphenic colouration (i.e., individuals vary markedly in their patterns and colours), they use adaptive colour change to match their backgrounds, microhabitat selection based on habitat colour and shape, and fast, intermittent motion (Bullough et al., 2023). Their extremely effective camouflage therefore presents challenges for visual detection, particularly as the human annotator will often fail to detect them until they are revealed by their movement. We used a video dataset of slaters filmed against their natural backgrounds around the coast in Falmouth (UK), and specified a primary static class (‘static’), and a primary motion class (‘moving’). Training used annotations from 130 frames, with 38 for validation from 12 videos. Annotations were only included if the slater was clearly visible to the annotator, and this decision was applied independently to the visual and static streams; slaters that were small, perfectly matched to their surrounds, or out of focus were typically easier to detect in the motion stream, and were not included in the static. We avoided repeatedly entering the same individual slater (and over-fitting the model for specific individuals) by using grey boxes to conceal them in subsequent frames unless they moved (see below). YOLO11s motion and static models were trained for 200 epochs. The static model achieved precision 0.875 and recall 0.846 (fig. S5). For 125 correct classifications, there were 23 false positives and 11 false negatives. The motion model had precision 0.834 and recall 0.922 (fig. S6). For 96 correct classifications there were 15 false negatives and 7 false positives.

The case studies here all use a default allocation of annotated frames to either training or validation datasets i.e. each annotated frame was given a 0.2 probability of being sent to the validation set. As such, each validation frame is likely to be from a video that was also included in the training dataset (with shared object and scene appearance), making them non-independent. We therefore ran a second test with validation and training annotations taken from mutually exclusive videos. Model outcomes were very similar with precision and recall for static of 0.908 and 0.859 (fig. S7); and motion of 0.893 and 0.876 (fig. S8).

### Semaphore flies

Semaphore flies (*Poecilobothrus nobilitatus*) have complex courtship behaviours that present a challenge for tracking and classification, with fast-moving limbs (wings and legs are often a blur at typical filming speeds), and fast flight. This example highlights some of the more advanced options available in the BehaveAI framework when working with more complex scenarios. The video database consisted of wild flies, filmed at 60 fps using a Sony a6400 with 60mm prime macro lens. Moving flies were highly salient in the motion stream, and three primary motion classes were specified (walk, fly and display). When the flies were not moving they were difficult to spot in the motion stream, so we also added a primary static class (rest) to find them from the static stream. However, the static classifier would be able to see all the cases of flies walking, flying or displaying that are not classed as rest. This would likely confuse the static classifier because a walking fly looks a lot like a resting fly. So we added `motion_blocks_static = true` to hide all the instances of walking, flying or displaying flies from the static classifier.

We also wanted to determine the sex of each fly from its wing markings, so added male and female as secondary static classes. However, when displaying or in flight these wing markings were not visible, so we could tell the model to ignore running the secondary classifier for these cases (`ignore_secondary = display, fly`). Finally, flies will often be detected by both the motion and static classifier, but the motion classifier will be more reliable and make very few false positive errors, so we set this to be the dominant stream for detections (`dominant_source = motion`). This only affects the video output - data from both streams are saved in the output.

Annotations from 524 frames were used for training, and 129 for validation, from 19 videos. YOLO11s classifiers were trained for 150 epochs (YOLO11s-cls for secondary models). The primary motion classifier achieved precision and recall of 0.933 and 0.932 respectively (fig. S9a), while the static model achieved 0.985 and 0.997 respectively (fig. S9b).. Secondary models (determining sex from the static images) achieved a reported accuracy from YOLO of 1 (i.e. correctly categorised all 57 males and 58 females while at rest fig. S9c-d).

Sample videos also highlight the individual tracking efficacy, with the Kalman filter based tracker successfully tracking the flies for extended periods despite instances of individual paths crossing.

### Edge performance

We tested real-time (‘edge’) performance of the framework on a low-end device (Raspbbery Pi 5 running Debian Trixie Pi OS 64-bit, with wide-angle, infrared OV5647 camera). We trained a primary motion classifier to detect the behaviour of adult *Tenebrio molitor* beetles on an oatmeal substrate, as either move, bury, or interact. The video was processed at 640×480 pixels (the same scale as model training), and operated at 8.3-9.6 frames-per-second, with the size of drop being linked to the number of concurrent detections. The change in framerate was the same between a model with a single classifier (Beetle), or three classifiers (move, bury, interact - where two or more beetles where in contact with one another). The system saved both video files of the detections (with a three second buffer either side of the period of detection), as well as the.csv file of coordinates, classifiers, and confidence levels. The same code runs at ∼20.6-24.5 fps on a conventional laptop (Framework Laptop 13 with AMD Ryzen 7040 Series CPU, Ubuntu 24.04) without GPU acceleration.

## Methods

### Colour-From-Motion

The framework operates by splitting video input into two parallel processing streams: the ‘static’ stream treats each frame as a conventional still colour image that provides shape, colour and textural information. The ‘motion’ stream presents temporal information using false colours that can intuitively be interpreted by both humans and CNNs to see patterns of movement (including speed, acceleration, and direction) over short timescales.

There are a range of user-selected options for creating colour from motion that are specified in the ‘BehaveAI_settings.ini’ file. Two different user-selectable motion strategies are available; the ‘exponential’ method calculates the absolute difference between the current frame and the previous frames, exponentially smoothing over successive frames to show different temporal ranges in different colour channels. With this mode, a moving object creates a white ‘difference’ image that leaves behind a motion blur that fades from white to blue, to green, to red. Increasing the exponential smoothing weights allows this method to show events further back in time at no extra computational processing or memory costs when running the classifier because there is no need to re-load any previous frames. This mode is better able to convey changes in speed within each frame; accelerating objects will outpace their red tail, creating a blue-to-green streak, while deceleration will allow the red tail to catch up, creating yellow-to-red tails. A further option (‘chromatic tails only’) removes the initial white difference information, preserving more static luminance information. This mode preserves the greatest proportion of luminance information, so is likely to suit tasks where detailed texture or form is key.

The ‘sequential’ method uses discrete frames rather than exponential smoothing, with colours coding the differences between the previous three frames (white, blue, green and red going back through time respectively), and is suited to classifying movements over this short range of frames and preserving more spatial information from previous frames (e.g., rather than a smooth tail, a flying animal’s characteristic wing shapes will remain visible over all four frames). The motion false colour is then optionally blended with the luminance channel to provide greater context – combining motion information with static luminance. The exponential weightings, false-colour composition, and degree of luminance blending are all user-adjustable. Frame skipping can also be used to perform measurements over a larger number of frames (longer time-span) at no additional processing costs (e.g., suited to slow-moving objects whose behaviour is more apparent from faster video playback). Settings can be altered following annotation, and the motion annotation dataset rebuilt using the new settings. The framework automatically detects changes in these settings if a model has already been trained, and asks whether the user would like to re-build and re-train with the new settings.

### Classifier Structure

We use the latest generation of YOLO classifiers (Jocher et al., 2023) for rapid and effective detection and classification from static and/or motion video streams. The user can select a model version (e.g., YOLO 11 or 8), and model size; larger models can improve accuracy, but require more computational resources. The nano (‘n’) or small (‘s’) models are generally sufficient. The framework supports both parallel and hierarchical processing of the static and motion streams. ‘Primary’ classifiers search each entire static and/or motion frame for objects (including objects of multiple types or ‘classes’); objects detected by a primary classifier are then automatically cropped and sent to ‘secondary’ static and/or motion classifiers (if a hierarchical model is specified). This allows the tasks of detection and classification to be performed using whichever stream (motion or static) is most appropriate. Users can specify the model structure with primary and (optional) secondary classes, together with the parameters for annotation, motion processing, and Kalman filter tracking (below), in the ‘BehaveAI_settings.ini’ file. Examples of different model structures are provided in the software release and below.

### Individual Tracking

The classification and tracking code uses a Kalman filter (Welch & Bishop, 1995) to keep track of individuals, assigning each a speed and heading that can be tracked even when paths overlap. Instances of multiple-detections are dealt with by combining boxes based on centroid distances and degree of overlap, and the user can select the prioritisation method when assigning a class to combined detections (e.g. static dominant, motion dominant, or highest confidence), although both streams are saved in the output file.

### Annotation

The framework’s annotation tool provides a user-friendly interface with initial manual annotation and training followed by optional semi-supervised (‘auto-annotation’, fig. 7). Semi-supervised annotation allows the user to learn where the CNNs are making errors, correct them for re-training, and repeat the annotation-training cycle until the desired performance is achieved. Initial manual annotation involves selecting input videos and drawing boxes around examples of the object classes, as is standard within deep learning pipelines. The interface additionally allows the user to flip between ‘static’ and ‘motion’ streams, with buttons or hotkeys for assigning primary and secondary classes. Following an initial limited annotation (e.g. 50 to 100 annotations per class depending on variance), we recommend training the model(s) to assist with further labelling; i.e. simply run the classifier script and it will handle model training automatically, see below. The annotation script then uses the trained model(s) to predict annotations, allowing the human annotator to accept model recommendations if they are accurate, but critically can also note any false positives or false negatives that the models make and correct them. Running the training again will now re-train the initial model. This workflow should increase the quality of the annotation dataset and speed of annotation; focusing on borderline cases and error correction, and reducing the risk of over-fitting or annotating more than is required. We also include a function for inspecting the annotated dataset, allowing for modification and error fixing.

**Figure 7.**
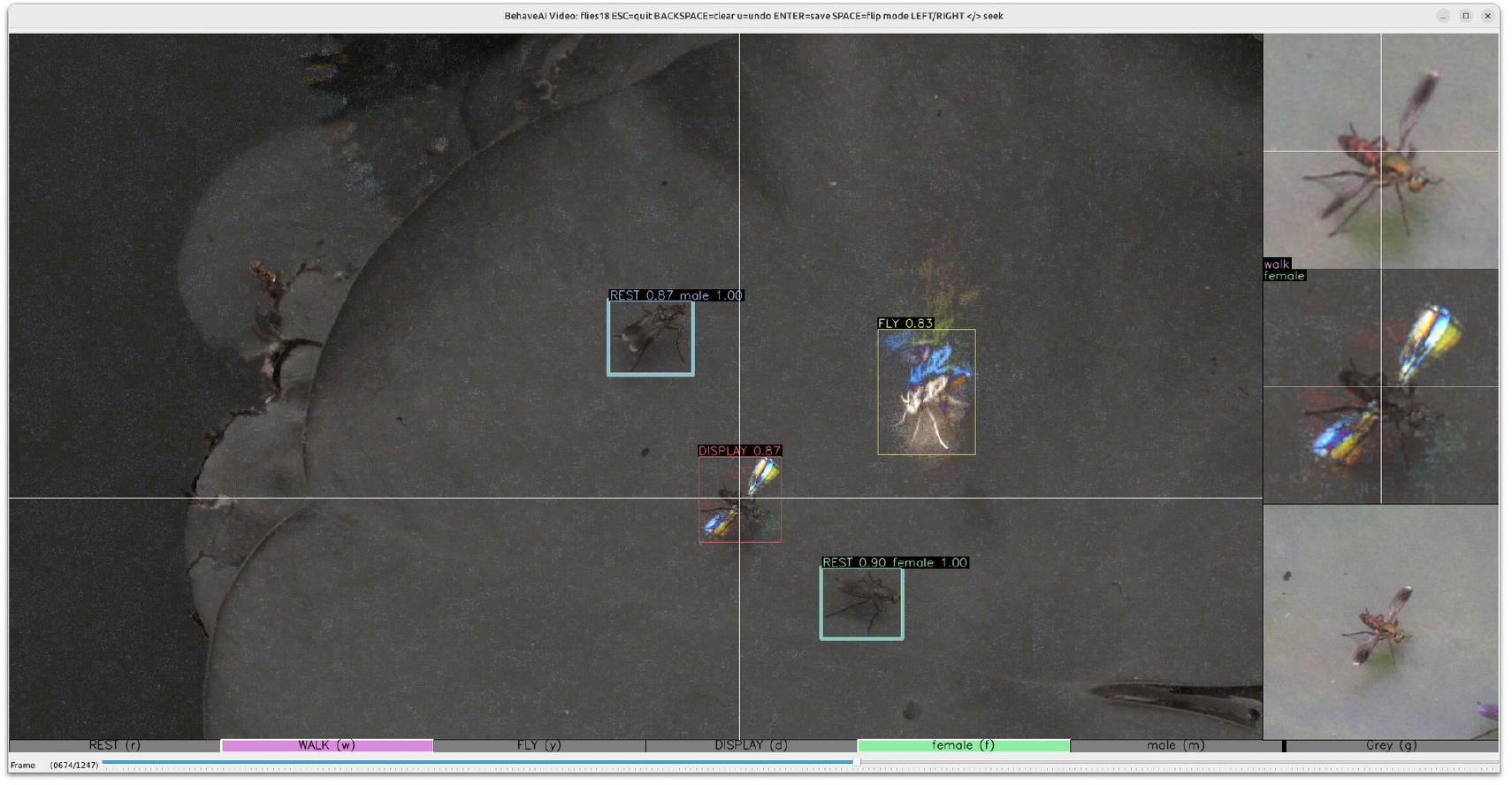
Annotation GUI screenshot. The main window shows the current motion frame (and can switch to static). The right-hand bar shows a zoom view of the current cursor position in static (top right) and motion streams (middle right), and also shows a looping animation of the video covering the same time-frame as the user-selected motion settings (bottom right). The bar at the bottom of the screen highlights the available primary (upper case) and secondary (lower case) classes, together with their associated keys. The track-bar at the bottom of the screen allows seeking through the video. Boxes in this example have all been drawn with auto-annotation, and show confidence levels for each (primary for behaviour and secondary for sex).

We have introduced a feature that optionally allows the user to shield parts of frames from the training and validation datasets by drawing a grey box over regions. Compared to a set of independent images, the frames of a video will often have near-identical objects and surrounds (e.g. in scenes where one target animal remains stationary while others are moving). Given all instances of a class present in any frame must be labelled for annotation, this would often require repeatedly labelling near-identical objects in successive frames. As such, models could easily become over-fitted for a specific instance, and lack the diversity of training and validation data to make robust inferences in other contexts. In these cases, drawing grey boxes over the target will shield them from the model and prevent over-fitting. Drawing grey boxes is also faster than precise bounding boxes, saving annotation time. There are also instances where the human annotator cannot be sure of the class (e.g. due to partial occlusion of an animal), in which case it would be potentially counterproductive to either label the object (and run the risk of the class being incorrect and limit model precision), or leave it unlabelled so that the model learns to avoid such instances (which could also be detrimental to model recall). Using a grey box can limit the dataset to only higher-confidence instances. Importantly, grey boxes should not be used for borderline cases where one animal behaviour or class is transitioning to another; it is important that the annotator uses a clear definition of the point at which these transitions occur and labels accordingly. Finally, grey boxes can also be used to shield moving instances from static frames, and/or static instances from motion frames. See the semaphore fly case study for an example where this is valuable.

### Batch Classification and Tracking

The classification and tracking tool manages both model training and batch input video processing, training initial models if none have been made. The tool also tracks the size of the annotation datasets and asks the user whether to re-train if additional annotations have been added. Any videos in the ‘input’ directory are batch processed, and the tool outputs labelled videos for inspection, together with a text (‘.csv.’) output listing each primary and (where relevant) secondary class for each individual, their centroid coordinates, plus confidence levels for each classifier.

### Case study processing

Processing was performed on a Framework 13 Laptop with Ubuntu 24.04, AMD Ryzen™ 7 7840U CPU (central processing unit), with NVIDIA GeForce RTX 2080 Ti, 11004MiB eGPU (external graphics processing unit, with CUDA support) unless otherwise specified.

### Raspberry Pi integration

We developed an installer and wrapper that allows for straightforward installation of the BehaveAI framework on a Raspberry Pi running the Raspbian operating system. Raspberry Pi-specific scripts also automatically handle the conversion of YOLO models to NCNN format, which makes more efficient use of the processing architecture available on Raspberry Pis.

## Author contributions

JT conceived and developed the initial tools, with wider support and project funding supported by KJG. Ongoing tool development and features were coded by JT with further testing, concepts and feature suggestions from TOW and JG. Raspberry Pi scripts were developed and tested by JG. The case studies were performed by JT. The GitHub user guide was written and maintained by JT and TOW. JT and TOW wrote the first draft, and all authors provided edits and feedback on subsequent drafts.

## Supplementary Figures

**Supplementary figure S1.**
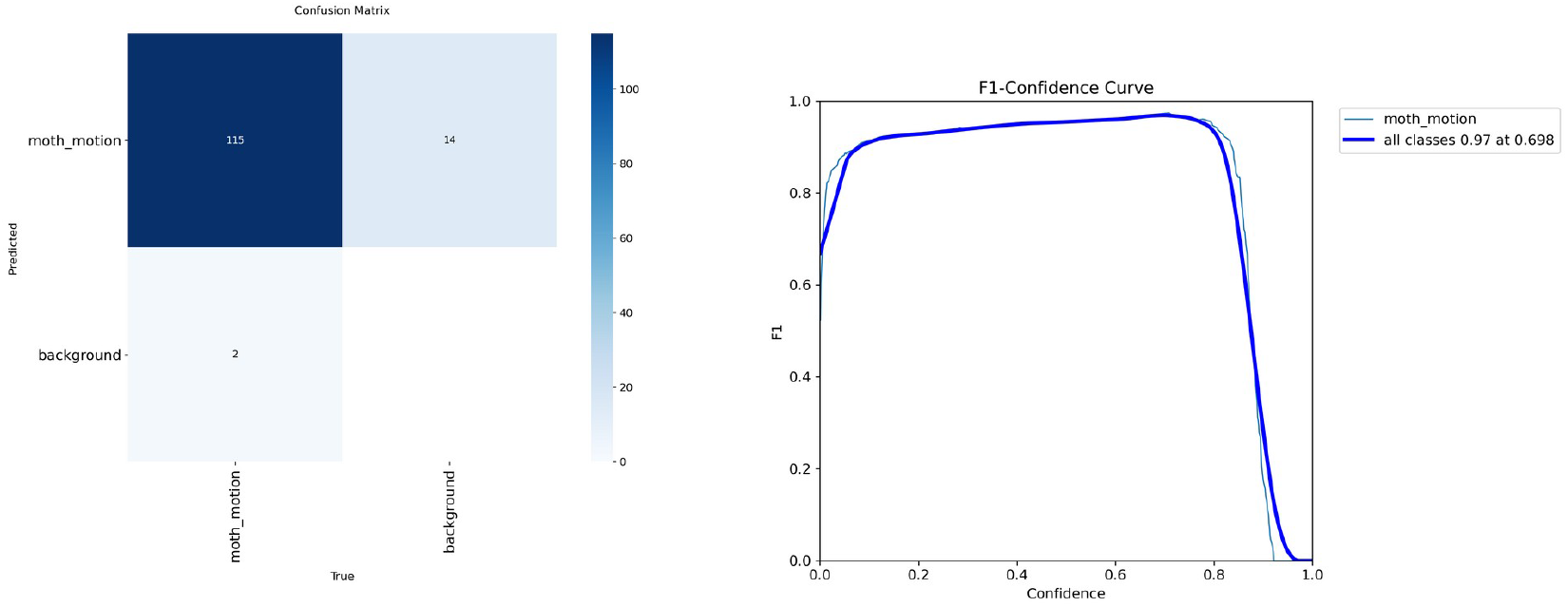
Confusion matrix and F1-confidence curves for grass moths classified from motion frames.

**Supplementary figure S2.**
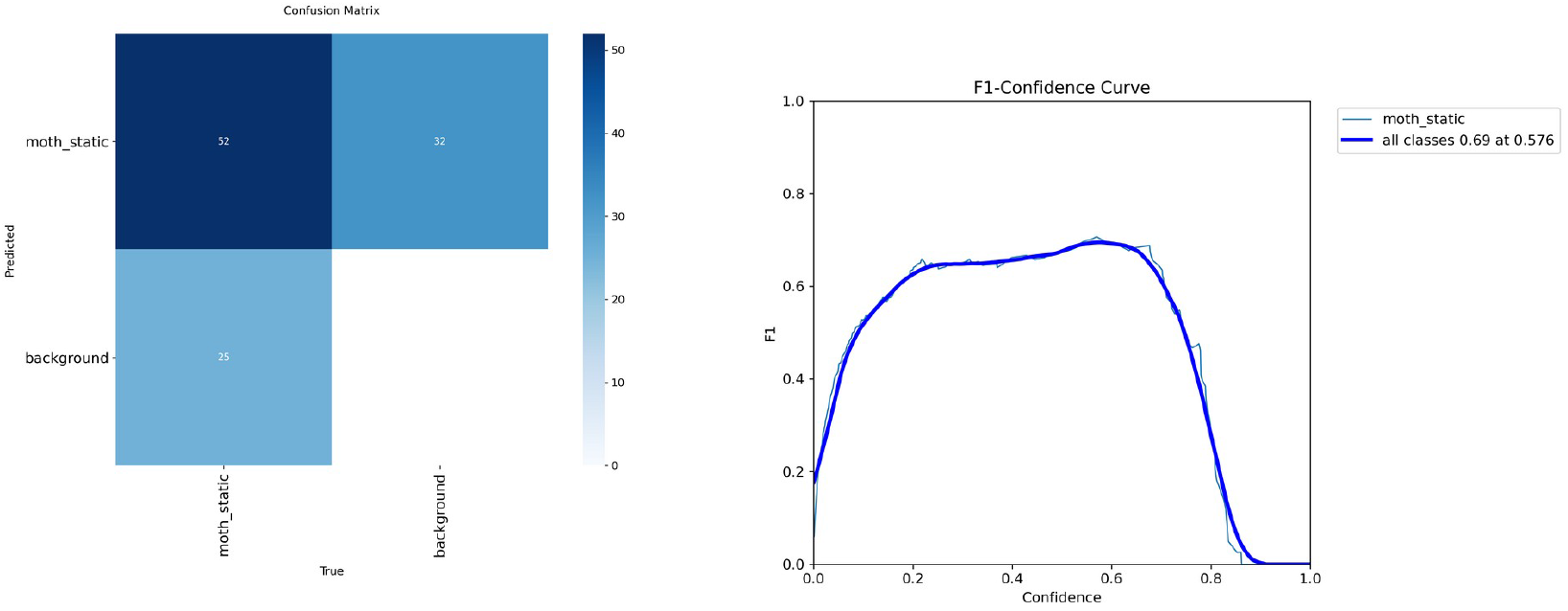
Confusion matrix and F1-confidence curves for grass moths classified from static frames.

**Supplementary figure S3.**
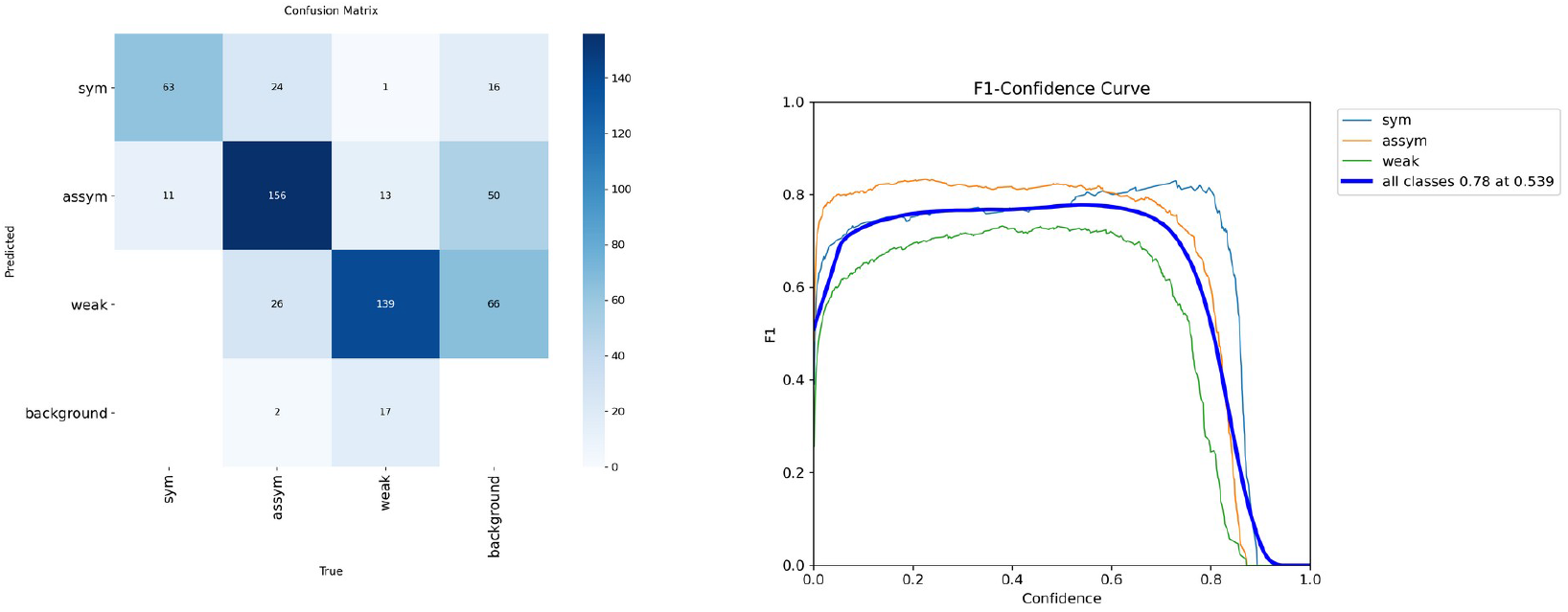
Confusion matrix and F1-confidence curves for sperm classified from motion frames.

**Supplementary figure S4.**
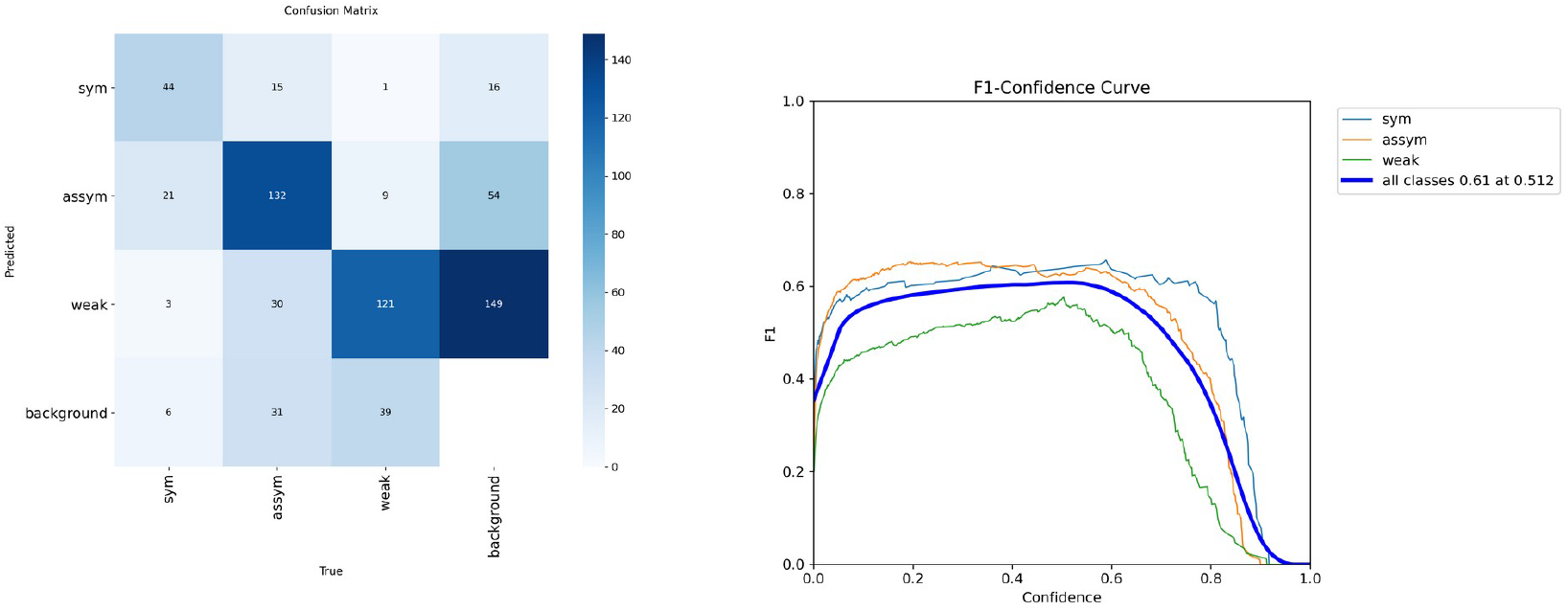
Confusion matrix and F1-confidence curves for sperm classified from static frames.

**Supplementary figure S5.**
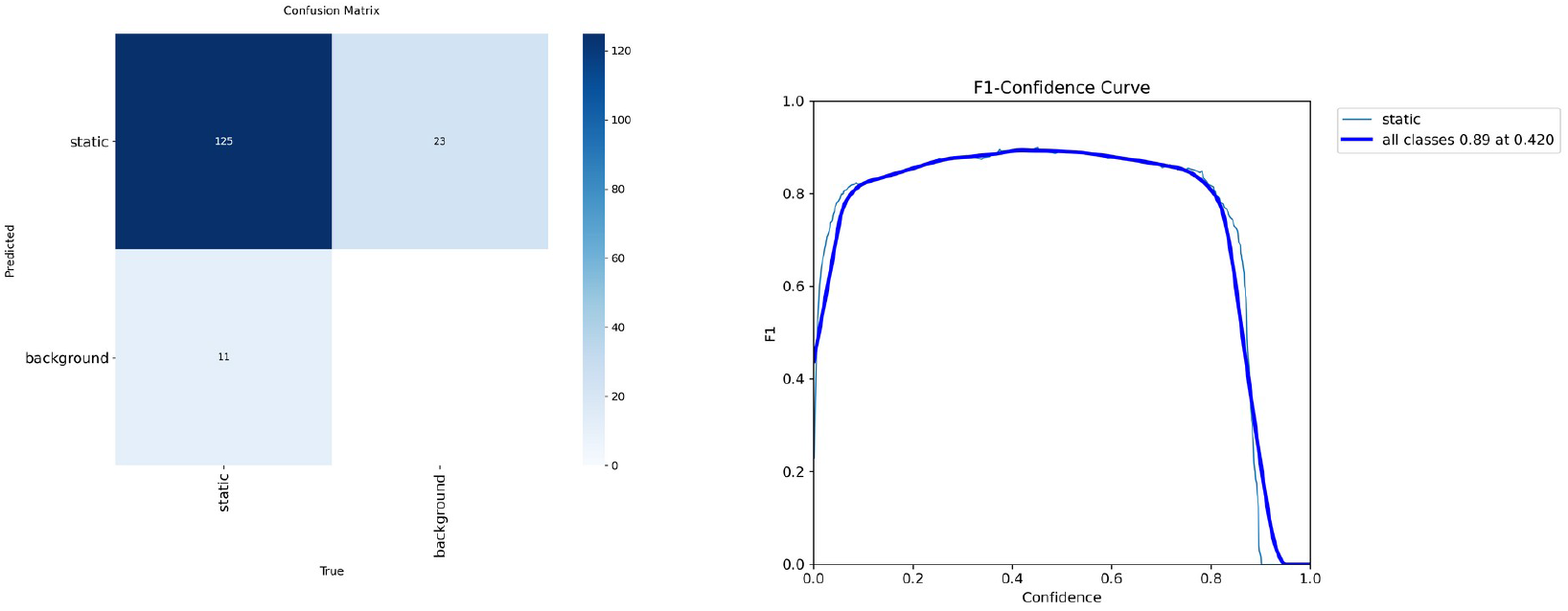
Confusion matrix and F1-confidence curves for sea slaters classified from static frames.

**Supplementary figure S6.**
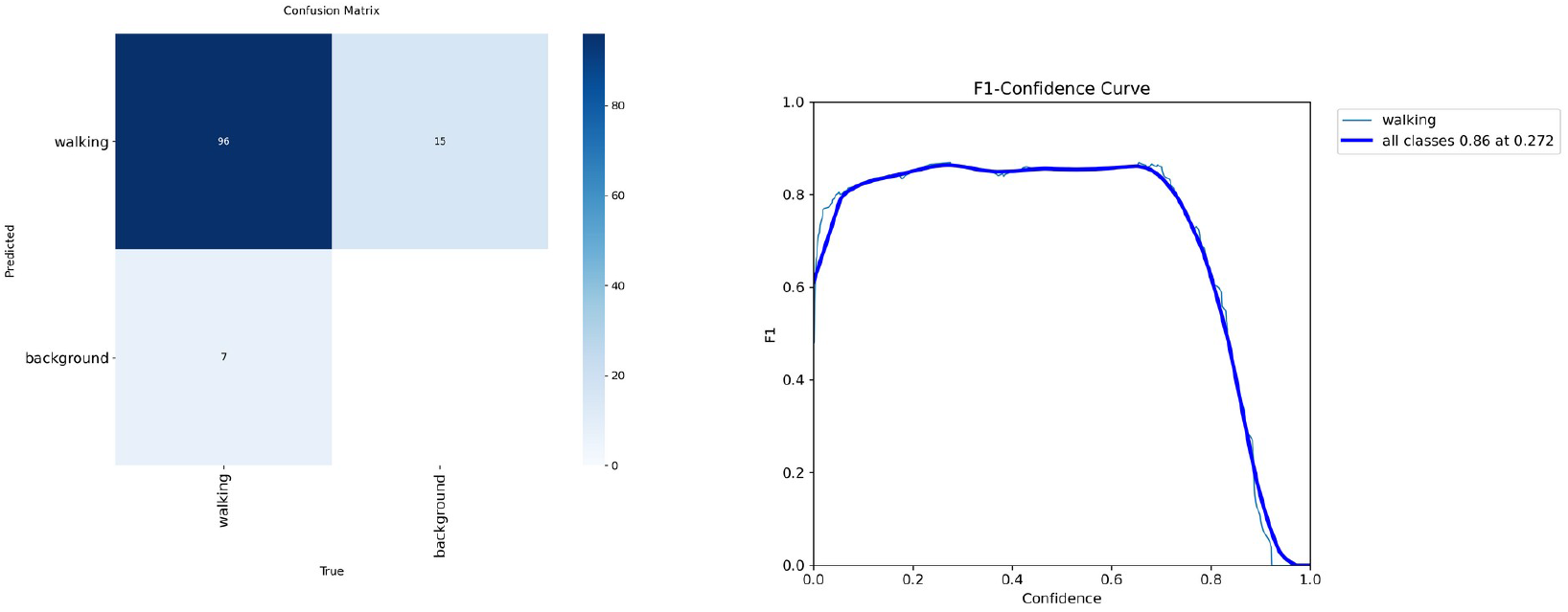
Confusion matrix and F1-confidence curves for sea slaters classified from motion frames.

**Supplementary figure S7.**
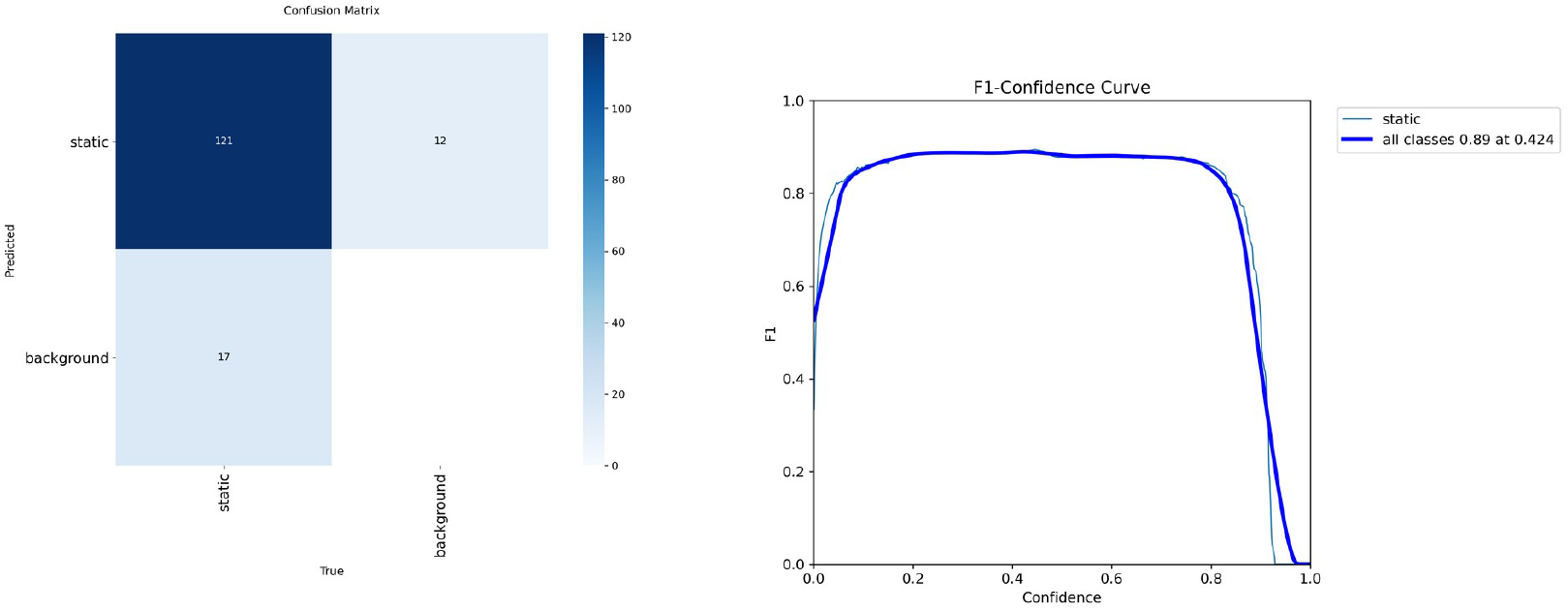
Confusion matrix and F1-confidence curves for sea slaters classified from static frames, with training and validation datasets taken from mutually exclusive videos.

**Supplementary figure S8.**
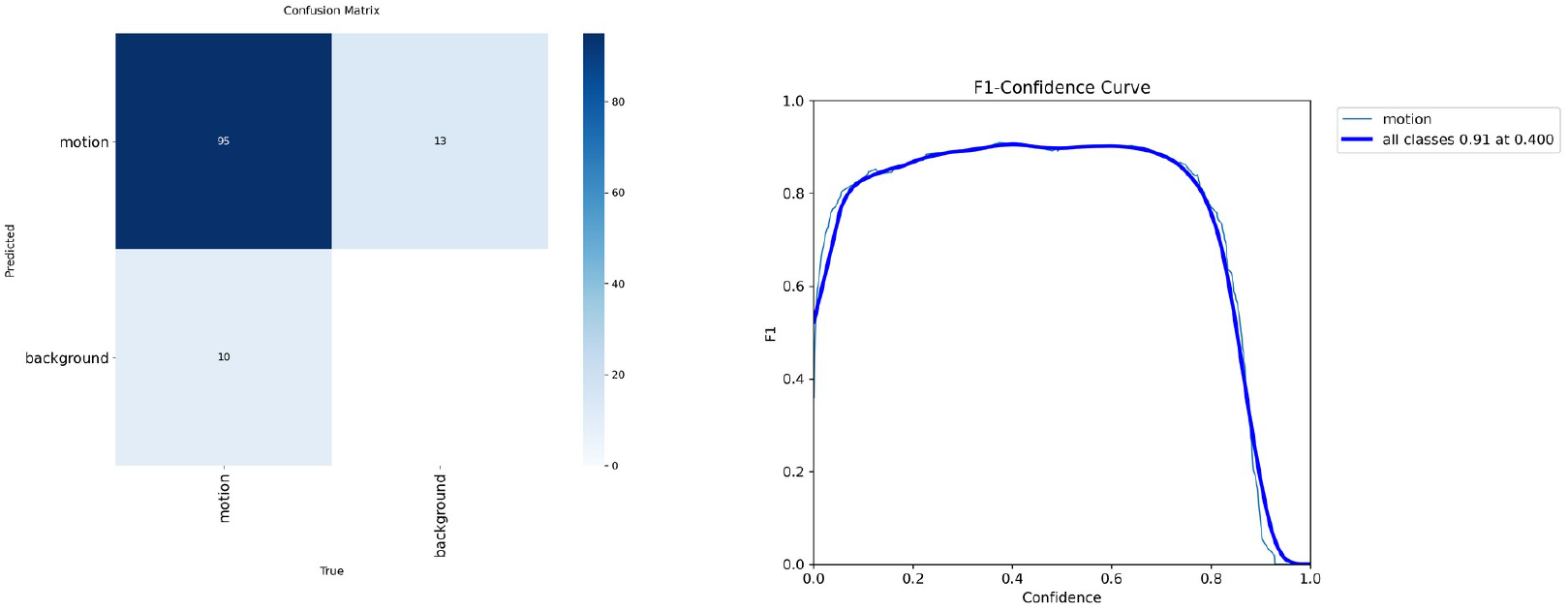
Confusion matrix and F1-confidence curves for sea slaters classified from motion frames, with training and validation datasets taken from mutually exclusive videos.

**Supplementary figure S9.**
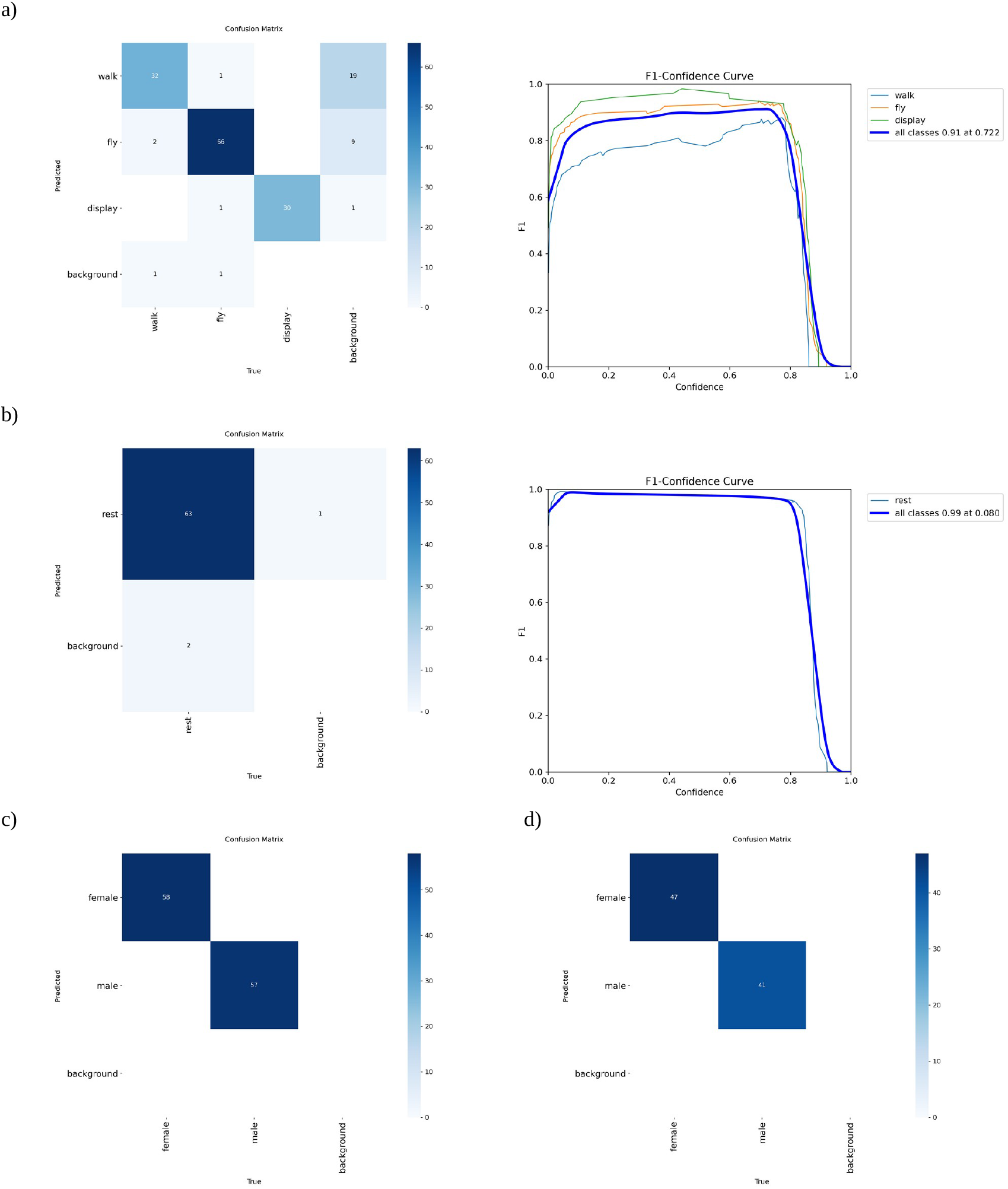
Confusion matrices and (for primary models) F1-confidence curves for semaphore flies. a) primary motion model, b) primary static model, c) secondary model determining sex while flies are at rest, and d) while flies are walking.

## References

Beery, S., Horn, G. van, & Perona, P. (2018). Recognition in Terra Incognita (No. arXiv:1807.04975). arXiv. 10.48550/arXiv.1807.04975

Bohnslav, J. P., Wimalasena, N. K., Clausing, K. J., Dai, Y. Y., Yarmolinsky, D. A., Cruz, T., Kashlan, A. D., Chiappe, M. E., Orefice, L. L., & Woolf, C. J. (2021). DeepEthogram, a machine learning pipeline for supervised behavior classification from raw pixels. Elife, 10, e63377.

Bouwmans, T. (2014). Traditional and recent approaches in background modeling for foreground detection: An overview. Computer Science Review, 11, 31–66.

Bullough, K., Gaston, K. J., & Troscianko, J. (2023). Artificial light at night causes conflicting behavioural and morphological defence responses in a marine isopod. Proceedings of the Royal Society B: Biological Sciences, 290(2000), 20230725. 10.1098/rspb.2023.0725

Carreira, J., & Zisserman, A. (2017). Quo vadis, action recognition? A new model and the kinetics dataset. Proceedings of the IEEE Conference on Computer Vision and Pattern Recognition, 6299–6308. http://openaccess.thecvf.com/content_cvpr_2017/html/Carreira_Quo_Vadis_Action_CVPR_2017_paper.html

Chan, A. H. H., Brookes, O., Waldmann, U., Naik, H., Couzin, I. D., Mirmehdi, M., Houa, N. A., Normand, E., Boesch, C., Boesch, L., Arandjelovic, M., Kühl, H., Burghardt, T., & Kano, F. (2025). Towards Application-Specific Evaluation of Vision Models: Case Studies in Ecology and Biology (No. arXiv:2505.02825). arXiv. 10.48550/arXiv.2505.02825

Chan, A. H. H., Putra, P., Schupp, H., Köchling, J., Straßheim, J., Renner, B., Schroeder, J., Pearse, W. D., Nakagawa, S., Burke, T., Griesser, M., Meltzer, A., Lubrano, S., & Kano, F. (2025). YOLO-Behaviour: A simple, flexible framework to automatically quantify animal behaviours from videos. Methods in Ecology and Evolution, 16(4), 760–774. 10.1111/2041-210X.14502

Hardin, A., & Schlupp, I. (2022). Using machine learning and DeepLabCut in animal behavior. Acta Ethologica, 25(3), 125–133. 10.1007/s10211-022-00397-y

Hu, Y., Ferrario, C. R., Maitland, A. D., Ionides, R. B., Ghimire, A., Watson, B., Iwasaki, K., White, H., Xi, Y., & Zhou, J. (2023). LabGym: Quantification of user-defined animal behaviors using learning-based holistic assessment. Cell Reports Methods, 3(3). https://www.cell.com/cell-reports-methods/fulltext/S2667-2375(23)00026-7

Jocher, G., Qiu, J., & Chaurasia, A. (2023). Ultralytics YOLO (Version 8.0.0) [Computer software]. https://ultralytics.com

Johansson, G. (1973). Visual perception of biological motion and a model for its analysis. Perception & Psychophysics, 14(2), 201–211. 10.3758/BF03212378

Komori, H., Isogawa, M., Mikami, D., Nagai, T., & Aoki, Y. (2023). Time-weighted motion history image for human activity classification in sports. Sports Engineering, 26(1), 45. 10.1007/s12283-023-00437-1

Krizhevsky, A., Sutskever, I., & Hinton, G. E. (2012). Imagenet classification with deep convolutional neural networks. Advances in Neural Information Processing Systems, 25. https://proceedings.neurips.cc/paper/2012/hash/c399862d3b9d6b76c8436e924a68c45b-Abstract.html

Mathis, A., Mamidanna, P., Cury, K. M., Abe, T., Murthy, V. N., Mathis, M. W., & Bethge, M. (2018). DeepLabCut: Markerless pose estimation of user-defined body parts with deep learning. Nature Neuroscience, 21(9), 1281–1289.

Milner, D., & Goodale, M. (2006). The visual brain in action (Vol. 27). Oup Oxford. https://books.google.co.uk/books?hl=en&lr=&id=8CaQDwAAQBAJ&oi=fnd&pg=PR7&dq=Milner+%26+Goodale+(1995/2006)+-+%22The+Visual+Brain+in+Action%22+(Oxford+Univ+Press)&ots=Pu4AbglDGa&sig=YM3CmAWyg0M_YeKvKOti8hdLVbA

Naik, H., Chan, A. H. H., Yang, J., Delacoux, M., Couzin, I. D., Kano, F., & Nagy, M. (2023). 3d-pop-an automated annotation approach to facilitate markerless 2d-3d tracking of freely moving birds with marker-based motion capture. Proceedings of the IEEE/CVF Conference on Computer Vision and Pattern Recognition, 21274–21284. https://openaccess.thecvf.com/content/CVPR2023/html/Naik_3D-POP_-_An_Automated_Annotation_Approach_to_Facilitate_Markerless_2D-3D_CVPR_2023_paper.html?utm_source=miragenews&utm_medium=miragenews&utm_campaign=news

Pereira, T. D., Tabris, N., Matsliah, A., Turner, D. M., Li, J., Ravindranath, S., Papadoyannis, E. S., Normand, E., Deutsch, D. S., & Wang, Z. Y. (2022). SLEAP: A deep learning system for multi-animal pose tracking. Nature Methods, 19(4), 486–495.

Raji, I. D., Bender, E. M., Paullada, A., Denton, E., & Hanna, A. (2021). AI and the Everything in the Whole Wide World Benchmark (No. arXiv:2111.15366). arXiv. 10.48550/arXiv.2111.15366

Ren, S., He, K., Girshick, R., & Sun, J. (2016). Faster R-CNN: Towards real-time object detection with region proposal networks. IEEE Transactions on Pattern Analysis and Machine Intelligence, 39(6), 1137–1149.

Simonyan, K., & Zisserman, A. (2014). Two-stream convolutional networks for action recognition in videos. Advances in Neural Information Processing Systems, 27. https://proceedings.neurips.cc/paper_files/paper/2014/hash/ca007296a63f7d1721a2399d56363022-Abstract.html

Thambawita, V., Hicks, S. A., Storås, A. M., Nguyen, T., Andersen, J. M., Witczak, O., Haugen, T. B., Hammer, H. L., Halvorsen, P., & Riegler, M. A. (2023). VISEM-Tracking, a human spermatozoa tracking dataset. Scientific Data, 10(1), 260. 10.1038/s41597-023-02173-4

Thapa, S., & Stachura, D. L. (2021). Deep Learning Approach for Quantification of Fluorescently Labeled Blood Cells in Danio rerio (Zebrafish). Bioinformatics and Biology Insights, 15, 11779322211037770. 10.1177/11779322211037770

Troscianko, T., Benton, C. P., Lovell, P. G., Tolhurst, D. J., & Pizlo, Z. (2009). Camouflage and visual perception. Philosophical Transactions of the Royal Society B: Biological Sciences, 364(1516), 449–461. 10.1098/rstb.2008.0218

Weinreb, C., Pearl, J. E., Lin, S., Osman, M. A. M., Zhang, L., Annapragada, S., Conlin, E., Hoffmann, R., Makowska, S., Gillis, W. F., Jay, M., Ye, S., Mathis, A., Mathis, M. W., Pereira, T., Linderman, S. W., & Datta, S. R. (2024). Keypoint-MoSeq: Parsing behavior by linking point tracking to pose dynamics. Nature Methods, 21(7), 1329–1339. 10.1038/s41592-024-02318-2

Welch, G., & Bishop, G. (1995). An introduction to the Kalman filter. https://gitea.auro.re/Krobot/krobotdocumentation/raw/commit/155a9eba0c1f173ff70f6e27361f642aa22cfb98/docs/doc/kalman.pdf

